# p53 binding sites in normal and cancer cells are characterized by distinct chromatin context

**DOI:** 10.1101/105221

**Authors:** Feifei Bao, Peter R. LoVerso, Jeffrey N. Fisk, Victor B. Zhurkin, Feng Cui

**Author notes:** These authors contributed equally to the paper as the first authors.

## Abstract

The tumor suppressor protein p53 interacts with DNA in a sequence-dependent manner. Thousands of p53 binding sites have been mapped genome-wide in normal and cancer cells. However, the way p53 selectively binds its cognate sites in different types of cells is not fully understood. Here, we performed a comprehensive analysis of 25 published p53 cistromes and identified 3,551 and 6,039 ‘high-confidence’ binding sites in normal and cancer cells, respectively. Our analysis revealed two distinct epigenetic features underlying p53-DNA interactions *in vivo*. First, p53 binding sites are associated with transcriptionally active histone marks (H3K4me3 and H3K36me3) in normal-cell chromatin, but with repressive histone marks (H3K27me3) in cancer-cell chromatin. Second, p53 binding sites in cancer cells are characterized by a lower level of DNA methylation than their counterparts in normal cells, probably related to global hypomethylation in cancers. Intriguingly, regardless of the cell type, p53 sites are highly enriched in the endogenous retroviral elements of the ERV1 family, highlighting the importance of this repeat family in shaping the transcriptional network of p53. Moreover, the p53 sites exhibit an unusual combination of chromatin patterns: high nucleosome occupancy and, at the same time, high sensitivity to DNase I. Our results suggest that p53 can access its target sites in a chromatin environment that is non-permissive to most DNA-binding transcription factors, which may allow p53 to act as a pioneer transcription factor in the context of chromatin.

## Introduction

The tumor suppressor p53 is a DNA-binding transcription factor (TF) that plays a pivotal role in preventing cancer growth (Levine 1997). p53 interacts with target DNA sites in a sequence-dependent manner, with the consensus motifs comprising two decanucleotides RRRCWWGYYY (R=A, G; Y=C, T; W=A, T), separated by a variable spacer (el-Deiry et al. 1992; Riley et al. 2008). Two approaches have been used to map p53 binding sites. The first approach is a traditional, single-gene method that usually uses reporter-gene constructs to determine the specific DNA fragments interacting with p53 *in vitro* (Riley et al. 2008), and these p53 sites are referred to as response elements (REs). The second approach is based on chromatin immune-precipitation (ChIP), an experimental technique used to investigate protein-DNA interactions *in vivo* (Johnson et al. 2007); these p53 sites are denoted as ChIP fragments. p53 REs and ChIP fragments are distributed differently in the human genome. Compared to REs, most p53 ChIP fragments are found far from transcriptional start sites (TSSs) (Wei et al. 2006), and some of them reside in interspersed repeats such as retroviral terminal repeats (LTRs) (Wang et al. 2007), short interspersed nuclear elements (SINEs) (Zemojtel et al. 2009; Cui et al. 2011) and long interspersed nuclear elements (LINEs) (Harris et al. 2009). Interestingly, quite a few p53 REs (Cui et al. 2011) and ChIP fragments (Zemojtel et al. 2009) reside in primate-specific Alu repeats, suggesting that Alu repeats play an important role in shaping the p53 regulatory network in the context of chromatin.

P53-DNA binding *in vivo* is influenced by local chromatin states (Beckeman and Prives 2010). The first evidence of this was reported in the seminal study by Espinosa and Emerson, who assessed the relative affinities of p53 for its two target sites within the *CDKN1A/p21* promoter and found that p53 exhibits a higher affinity to the sites in the context of chromatin than in naked DNA (Espinosa and Emerson 2011). Moreover, p53-dependent changes in histone acetylation have been observed at the promoters of many p53 target genes, including *CDKN1A/p21* (Barlev et al. 2011) and *MDM2* (Candau et al. 1997). H3 and H4 histone modifications have been observed for 20% of target genes after p53 is overexpressed (Vrba et al. 2008; Kaneshiro et al. 2007). Recent studies revealed that the transcriptionally active histone mark, histone H3 lysine 4 trimethylation (H3K4me3), is associated with both p53 binding and transactivation of target genes (Menendez et al. 2013), and the repressive histone mark, H3K27me3, is detected in repressed p53 target genes (Li et al. 2012). All these observations suggest that p53 binding coordinates with histone modifications to regulate gene transcription. The level of DNA methylation at p53 binding sites was also shown to be critical for *in vivo* DNA binding (Botcheva et al. 2011).

TF-DNA binding *in vivo* is usually associated with open chromatin, which is characterized by increased DNase I sensitivity (Boyle et al. 2008), and an integrative analysis of ChIP fragments bound by 119 human TFs revealed that most TF binding sites are located in nucleosome-depleted DNase I sensitive regions (Wang et al. 2012). However, another group of proteins (including p53) known as pioneer factors are able to interact with nucleosomal DNA (Zaret and Carroll 2011; Iwafuchi-Doi and Zaret 2014; Zaret and Mango 2016). Indeed, it has been shown that p53 can interact with nucleosomal DNA both *in vitro* (Sahu et al. 2010; Laptenko et al. 2011) and *in vivo* (Laptenko et al. 2011; Lidor et al. 2010). This finding is consistent with previous studies showing that well-known p53 REs including the *p21* 5’ RE are often exposed on the nucleosomal surface (Cui and Zhurkin 2014) and occur in chromatin domains that are resistant to DNase I digestion (Braastad et al. 2003). Recent studies however have showed that the binding sites of two pioneer factors, progesterone receptor (PR) and forehead box protein A2 (Foxa2), occur in genomic regions with both high nucleosome occupancy and high sensitivity to DNase I (Ballare et al. 2013a; Ballare et al. 2013b; Iwafuchi-Doi et al. 2016), suggesting that high sensitivity to DNase I does not necessarily reflect nucleosome depletion. Whether p53 *in vivo* binding sites also occur in genomic regions displaying high DNase I sensitivity remains unknown.

All these studies suggest that p53-DNA binding *in vivo* is associated with multiple chromatin patterns (i.e., histone modifications, DNA methylation, nucleosome occupancy and DNase I sensitivity). Because normal cells (NCs) and cancer cells (CCs) are characterized by drastically different chromatin organization (Jones and Baylin 2007; Baylin and Jones 2011), we hypothesize that p53 binding sites derived from the two different types of cells exhibit distinct epigenetic patterns.

In the present study, we conducted a comprehensive analysis of 25 published p53 cistromes that include more than 120,000 p53 ChIP fragments from numerous NC and CC lines under various stress conditions. Merging these datasets produces 6,039 NC and 3,551 CC ‘high-confidence’ p53 ChIP clusters that include three or more overlapping ChIP fragments. These clusters are enriched in the endogenous retrovirus 1 (ERV1) repeat family of the human genome. We found that regardless of the cell type, the p53 ChIP clusters occur in genomic regions with high nucleosome occupancy and high DNase I sensitivity, which differs from most DNA-binding TFs. Finally, compared to their counterparts in cancer cells, the NC ChIP clusters are more likely associated with a higher level of transcriptionally active histone marks (H3K4me3 and H3K36me3) and DNA methylation, but a lower level of repressive histone marks (H3K27me3). In light of these findings, a new scheme was proposed to explain the distinct p53 genomic binding patterns in normal and cancer cells.

## Results

### Comparison between p53 binding sites identified *in vitro* and *in vivo*

To compare the p53 binding sites identified *in vitro* and *in vivo*, we collected 154 p53 REs (Supplemental Table S1) and ~120,000 p53 ChIP fragments from 25 published datasets, which include 42,020 NC ChIP fragments in 9 datasets and 77,796 CC ChIP fragments in 16 datasets (Fig. 1A, Supplemental Table S2). The NC ChIP fragments are significantly longer than those from CC lines (Wilcoxon test, *p* < 2.2 × 10^−16^) (Supplemental Fig. S1). Note that the longest p53 ChIP fragments came from human embryonic stem cells (hESCs) (Akdemir et al. 2014) and keratinocytes (McDade et al. 2014) (Supplemental Fig. S2). By contrast, the CC ChIP fragments have a narrower length distribution, with an average length of approximately 500 bp (Supplemental Fig. S3). Although the possibility of experimental variations cannot be ruled out, this difference in ChIP fragment lengths could be due to distinct chromatin structures in NCs and CCs. That is, cancer chromatin may have a more ‘extended’ structure due to global hypomethylation (Jones and Baylin 2007; Baylin and Jones 2011), which would facilitate chromatin shearing in ChIP-seq experiments.

**Fig. 1.**
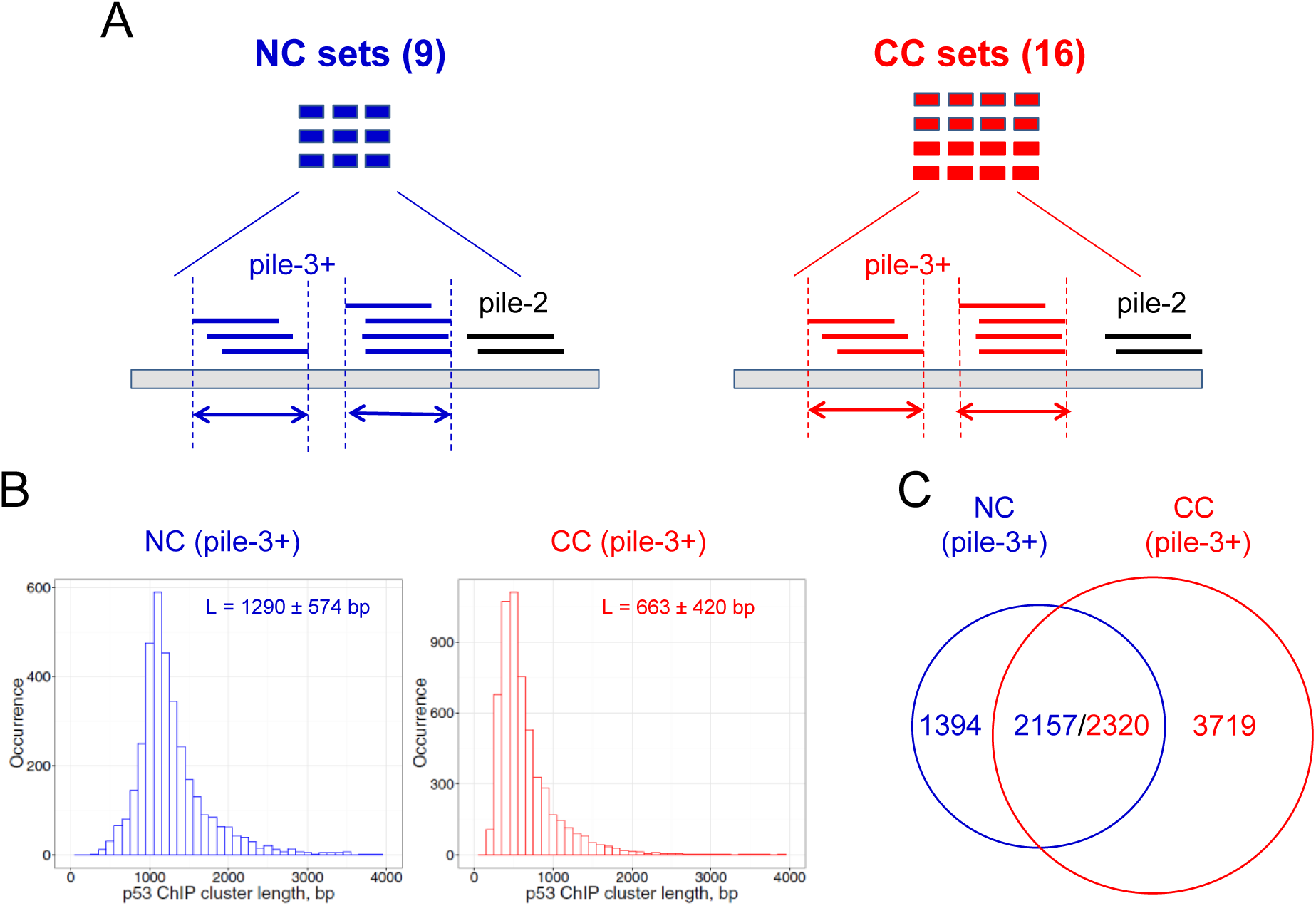
Analysis of p53 ChIP clusters derived from human normal and cancer cells. (A) Schematic view of ChIP fragment analysis. Nine datasets from normal cell (NC) lines and 16 datasets from cancer cell (CC) lines were used in this study (Supplemental Table S2). The positions of ChIP fragments were converted to human genome assembly hg18 using the LiftOver utility in the UCSC Genome Browser. Overlapping ChIP fragments form clusters. A pile-3+ cluster containing three or more overlapping members is denoted as a ChIP cluster and used for further analysis. Arrows denote the lengths of clusters. (B) Length distribution of ChIP clusters from NC (left) and CC (right) datasets. The x-axis represents ChIP cluster lengths binned into 100-bp intervals. The y-axis represents the occurrence of ChIP clusters in each bin. (C) Overlap of ChIP clusters from NCs and CCs. The p53 ChIP clusters are naturally divided into three groups: NC-only, N/C, and CC-only.

To investigate how p53 REs identified *in vitro* overlap with p53 ChIP fragments obtained *in vivo*, we visualized p53 ChIP fragments in known target genes using the UCSC Genome Browser (Supplemental Fig. S4). We found that out of 154 p53 REs, 71 REs (46%) overlap with at least one NC or CC ChIP fragment (Supplemental Table S1), consistent with earlier studies (Botcheva et al. 2011).

For a detailed comparison, we selected two well studied p53 target genes, *CDKN1A/p21* (el-Deiry et al. 1993) and *PUMA* (Yu et al. 2001). We found that multiple ChIP fragments contain previously reported p53 REs in these two genes (Supplemental Fig. S5 and S6), demonstrating a good agreement between p53 REs and ChIP fragments. Analysis of ChIP fragments in these two genes led to the following observations.

First, in addition to known p53 REs, ChIP fragments overlap at other locations, which may have functional significance. For example, in the *p21* gene, multiple ChIP fragments are located at an intronic locus, ~3.3 kb downstream of TSS (black arrow in Supplemental Fig. S5). This genomic locus corresponds to a previously identified p53 binding site that can activate a variant of the *p21* gene, *p21B*, to induce apoptosis (Nozell and Chen 2002). This result suggests that superimposition of multiple ChIP fragments from different datasets helps identify functionally important p53 binding sites.

Second, normal and cancer cells differ in p53 ChIP occupancy at previously reported p53 REs. Consistent p53 occupancy is often observed in cancer but not in normal cell lines. For example, 14 out of 16 (88%) CC ChIP datasets show p53 occupancy at the 5’ or 3’ REs in the *p21* promoter, whereas only 4 out of 9 (44%) NC datasets have signals at these positions (Supplemental Fig. S5). Likewise, 12 out of 16 (75%) CC datasets have p53 occupancy at the known *PUMA* RE, whereas only 2 out of 9 (22%) NC datasets show signals around the same position (Supplemental Fig. S6). Overall, we observed distinctive p53 binding patterns in normal and cancer cells, in agreement with earlier studies (Shaked et al. 2008; Botcheva et al. 2011; Botcheva et al. 2014).

### Searching for high-confidence p53 binding sites in normal and cancer cells

We set out to assemble the NC and CC ChIP fragments into clusters across the human genome (hg18). About 40% of the ChIP fragments are singletons that are scattered throughout the genome, whereas the remaining fragments overlap with each other and form clusters (Fig. 1A and Supplemental Table S3). The clusters are denoted pile-2, pile-3, pile-4, and so forth based on their coverage (i.e., number of overlapping members in a cluster). Our Monte Carlo simulation analysis showed that pile–3+ clusters are highly unlikely to form by chance (*p* < 0.01, see Methods and Supplemental Table S3), which suggests that these clusters reflect ‘real’ ChIP enrichment in normal and cancer cells.

Thus, we identified 3,551 NC and 6,039 CC pile-3+ clusters (for brevity, denoted NC and CC ChIP clusters, respectively). We found that the lengths of NC and CC ChIP clusters differ significantly (Wilcoxon test, *p* < 2.2 × 10^−16^, Fig. 1B). We also found that 2,157 (61%) of the NC ChIP clusters overlap with the CC ChIP clusters, whereas 2,320 (38%) of the CC ChIP clusters overlap with the NC ChIP clusters (Fig. 1C, Supplementary Tables S4-S5). This small discrepancy in the number of overlapping clusters (2,157 vs. 2,320) is likely due to the different length distributions in NC and CC ChIP clusters (Fig. 1B). In other words, a typical NC ChIP cluster (~1,300 bp in length) may cover two CC clusters (~700 bp in length). The 2,320 ChIP clusters from CCs were included in the “normal/cancer” (*i.e.*, N/C) group for the studies described below (because they have a higher number than their counterparts from normal cells). The ChIP clusters that are not shared by the normal and cancer datasets are put into the “CC-only” group or the “NC-only” group, respectively.

Noticeably, most p53 ChIP clusters occur more than 10 kb away from the TSSs of the nearest genes, which is in sharp contrast to p53 REs (Chi-squared test *p* < 6.9 × 10^−7^, Supplemental Fig. S7). This result indicates that p53 binding sites identified *in vitro* and *in vivo* exhibit distinctive genomic distributions, consistent with earlier studies (Wei et al. 2006; Cui et al. 2011).

Gene ontology analysis by DAVID (Huang et al. 2009a; Huang et al. 2009b) revealed that ChIP clusters are enriched in genes involved in p53 signaling pathways. Consider the N/C group as an example. The most significant KEGG pathway is the “p53 signaling pathway”, which has an enrichment score of 9.6 (Fisher’s exact test, *p* < 0.01) (Supplemental Fig. S8A). For this reason, most of enriched gene ontology terms are related to p53 functions, including “regulation of apoptosis”, “DNA damage response”, and “regulation of cell cycle” (Supplementary Fig. S8B).

### Distribution of p53 motifs in high-confidence binding sites in human repetitive elements

To examine the relationship between p53 ChIP clusters and repetitive elements in the human genome, we first identified p53 motifs (~20 bp in length) in each cluster using our previously developed computational method (Cui et al. 2011) based on a position weight matrix (PWM) formalism (Figure 2A). This method scores DNA fragments based on their similarity to the consensus p53 binding motif. That is, the more similar to the consensus sequence, the higher the PWM score. A cutoff of 70% suggested in our previous study (Cui et al. 2011) was used. If multiple p53 motifs with the score 70% or higher are identified in a given p53 ChIP cluster, the motif with the highest score is selected. As a result, 3,298 (93%) motifs were identified in 3,551 NC ChIP clusters, whereas 4,791 (76%) motifs were found in 6,309 CC ChIP clusters (Supplemental Table S4-S5). Note that a substantial fraction of p53 ChIP clusters (~7% NC and ~24% CC clusters) contains no obvious p53 binding motifs; a similar trend was also observed in other studies (Wei et al. 2006; Smeenk et al. 2008).

**Fig. 2.**
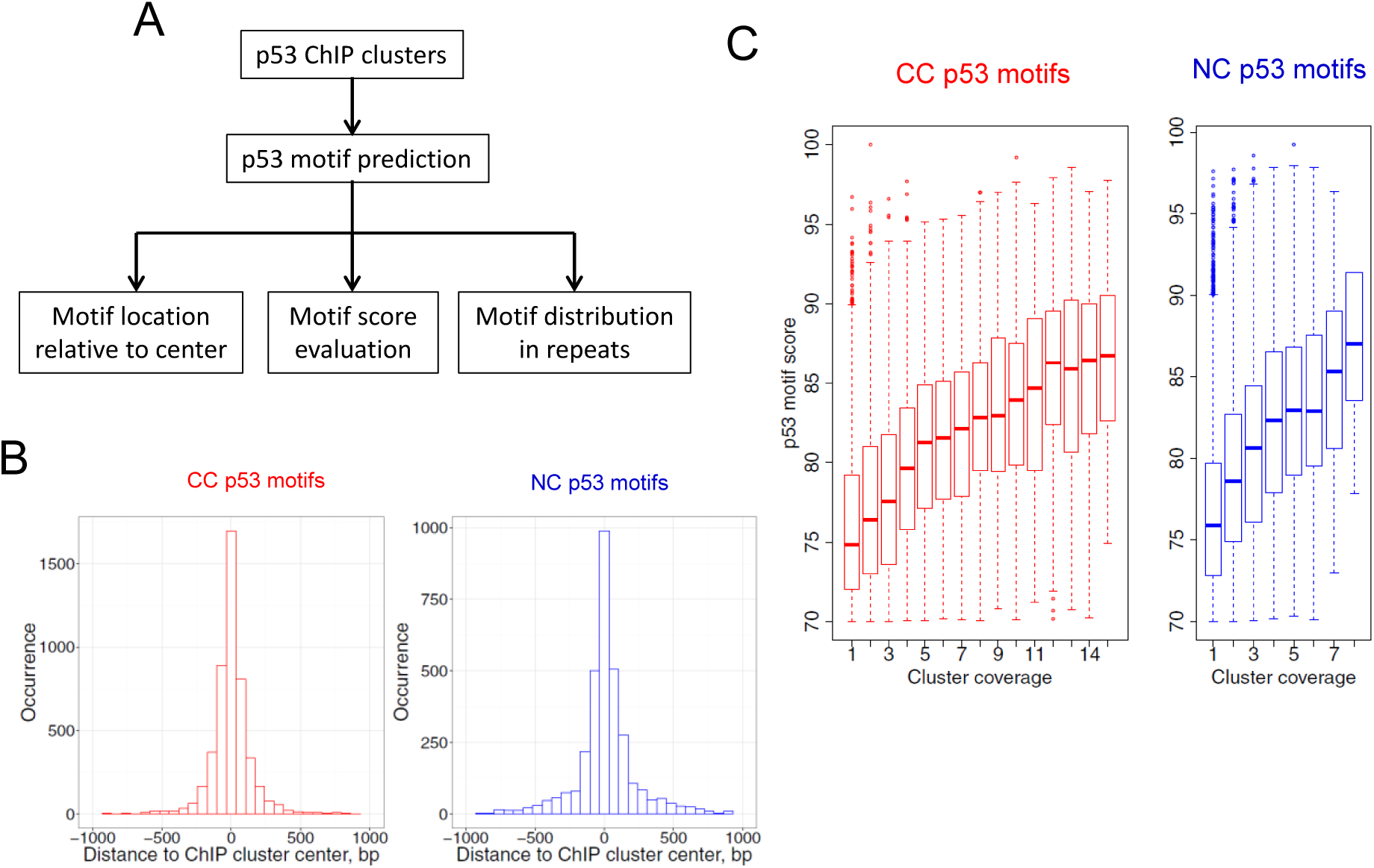
Analysis of p53 motifs in ChIP clusters. (A) Overall research plan for the p53 motifs identified in ChIP clusters. The PWM-based tool developed in our previous study (Cui et al. 2011) was used to predict p53 motifs. If multiple p53 motifs were found in a given cluster, the motif with the highest PWM score was selected. (B) Locations of p53 motifs relative to the ChIP cluster centers. (C) Distribution of p53 motif scores with respect to the coverage of ChIP clusters.

There are several possible reasons for the lack of p53 binding motifs in these ChIP clusters. First, in certain cases, p53 binds DNA based on the topology of a DNA fragment, not on its sequence (Gohler et al. 2002). Second, p53 can bind to different motifs such as microsatellites in the *PIG3* gene (Contente et al. 2002). Third, p53 can be recruited to target sites through indirect DNA binding, as shown in the p53-dependent transcriptional repression of the *survivin* gene (Fischer et al. 2015).

We found that the identified p53 motifs tend to occur close to the centers of p53 ChIP clusters (Figure 2B). Moreover, the average motif scores increase with the coverage of p53 clusters (Figure 2C), indicating that the clusters with more overlapping members tend to have stronger p53 motifs. To check whether the p53 motifs overlap with repeat elements, we compared the locations of motifs and the RepeatMasker annotation of the human genome (hg18). We found that about 40% of the p53 motifs overlap with various types of repetitive elements, including LINE, SINE, LTR and simple repeats (Fig.3A-C), consistent with earlier studies (Wang et al. 2007; Zemojtel et al. 2009; Harris et al. 2009; Cui et al. 2011). Analysis of the repeat-associated p53 motifs yielded the following observations.

**Fig. 3.**
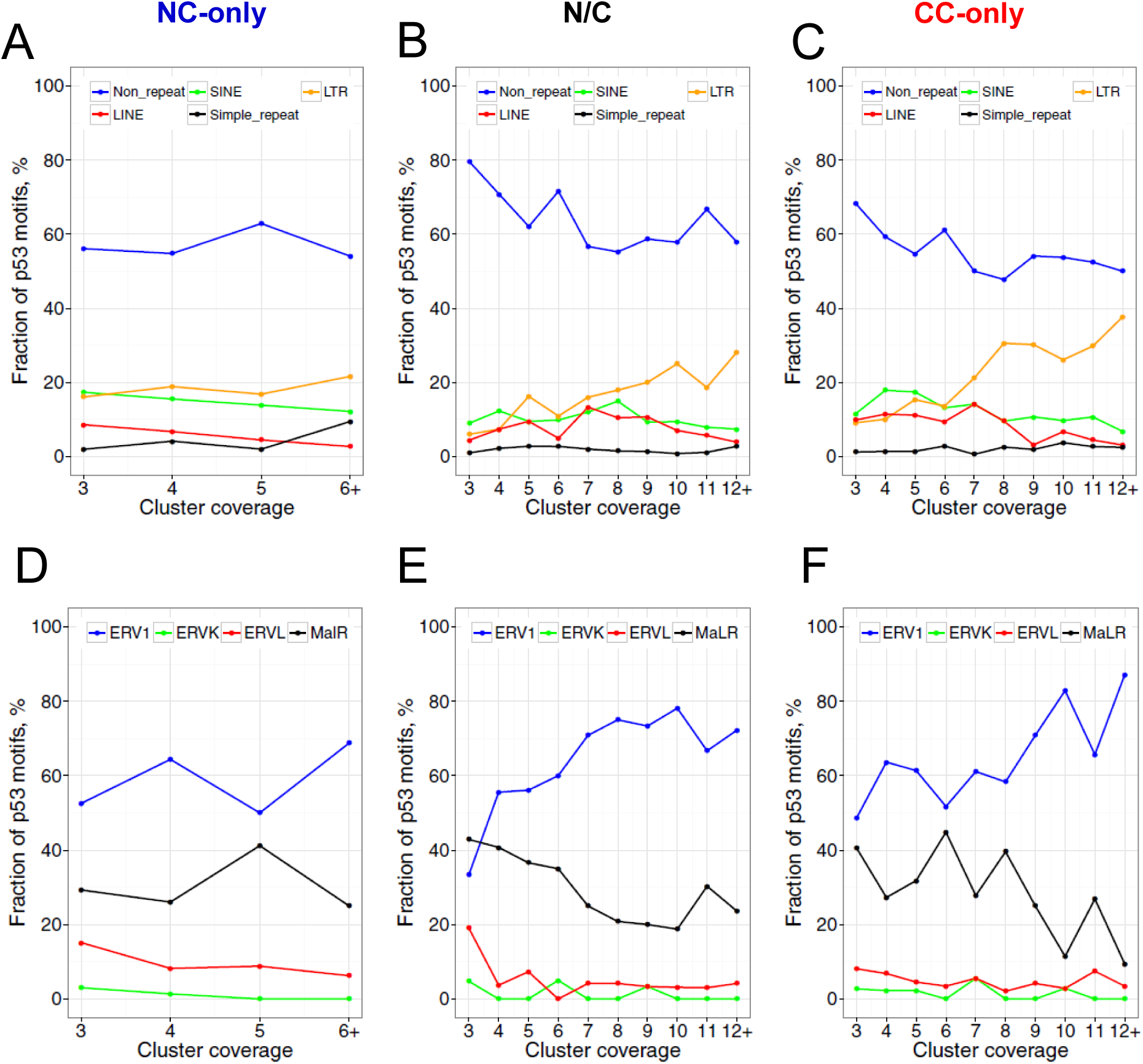
Distribution of identified p53 motifs in the human genome (A-C) and in the ERV/LTR repeat family (D-F). ChIP clusters from the NC-only group (A, D), the N/C group (B, E) and the CC-only group (C, F) were taken into consideration. RepeatMasker annotation of the human genome (hg18) downloaded from the UCSC Genome Browser was used for analysis. If a given p53 motif does not overlap with any repetitive element in the genome, it was designated as “Non_repeat”. Otherwise, the motif was assigned to the corresponding repeat family and subfamily. The fractions of p53 motifs in repeat (sub)families and in the “Non_repeat” group were calculated with respect to the coverage of ChIP clusters.

First, the fraction of repeat-associated p53 motifs increases with the coverage of the clusters (Fig. 3A-C). That is, the high-coverage p53 ChIP clusters tend to occur in repetitive regions. Because the high-coverage clusters are likely to have stronger p53 motifs (Fig. 2C), it follows that strong p53 motifs are likely to reside in repeats, consistent with earlier studies (Su et al. 2015). Second, the fraction of LTR-associated p53 motifs increases with the coverage of the clusters (orange lines in Fig. 3A-C), indicating that p53 motifs are usually located in the repeat elements of the LTR class. Third, the fraction of ERV1-associated p53 motifs increases with the coverage of the clusters (blue lines in Fig. 3D-F). Overall, about 12% of identified p53 motifs occur in ERV1 elements, which is much higher than the fraction of these elements (2.9%) in the human genome (Lander et al. 2001). The percentage distribution in repeat families (Fig.3A-C) and subfamilies (Fig.3D-F) is significantly different from the corresponding fractions in the human genome (Lander et al. 2001) for all groups and all coverage of clusters (Chi-squared test, *p* < 0.05), except for the pile-3 clusters in Figure 3E.

Overall, our data is generally consistent with the conclusion of previous studies (Wang et al. 2007), that p53 binding sites are highly enriched in ERV1 repeats. Moreover, we showed that ERV1 is the only repeat family in which NC and CC ChIP clusters are enriched, suggesting that other families such as SINE, LINE and simple repeats may play a less important role in p53 transcriptional programs.

### High nucleosome occupancy and DNase I sensitivity at high-confidence p53 binding sites

To examine the chromatin organization around p53 binding sites, we started with two well-studied p53 target sites, the 5’ RE and 3’ RE in the *CDKN1A/p21* promoter. It has been shown that both p21 REs are located within nucleosomal DNA *in vitro* and *in vivo* (Laptenko et al. 2011). In such a case, we would expect to see that both 5’ and 3’ REs occur in genomic regions with relatively high nucleosome occupancy in the chromatin landscape of normal (GM12878) and cancer (K562) cell lines. Indeed, we observed increased nucleosome occupancy around the two *p21* REs (Supplemental Fig. S9), which is about two times higher than the average occupancy in the human genome.

To check whether this tendency holds for the NC and CC p53 ChIP clusters, we calculated *in vivo* nucleosome occupancy profiles around the ChIP clusters in the NC-only, N/C and CC-only groups. Clearly, for all three groups, the centers of ChIP clusters are located in the genomic regions with a significantly higher nucleosome occupancy compared to flanking DNA 1000 bp away (Fig. 4A-C, Wilcoxon test, *p* < 0.01). To determine whether the observed high nucleosome occupancy is related to intrinsic histone-DNA interaction, we used published data on the positioning of reconstituted human nucleosomes (Valouev et al. 2011) and constructed *in vitro* nucleosome occupancy profiles for the p53 ChIP clusters. The nucleosome occupancy at the cluster centers is much higher compared to flanking DNA (Supplemental Fig. S10) and this difference is statistically significant (Wilcoxon test, *p* < 10^−15^, Supplemental Table S6). Thus, our results are consistent with the earlier findings that p53 binding sites usually reside within genomic regions with DNA sequences predicted to encode relatively high intrinsic nucleosome occupancy (Lidor et al. 2010).

**Fig. 4.**
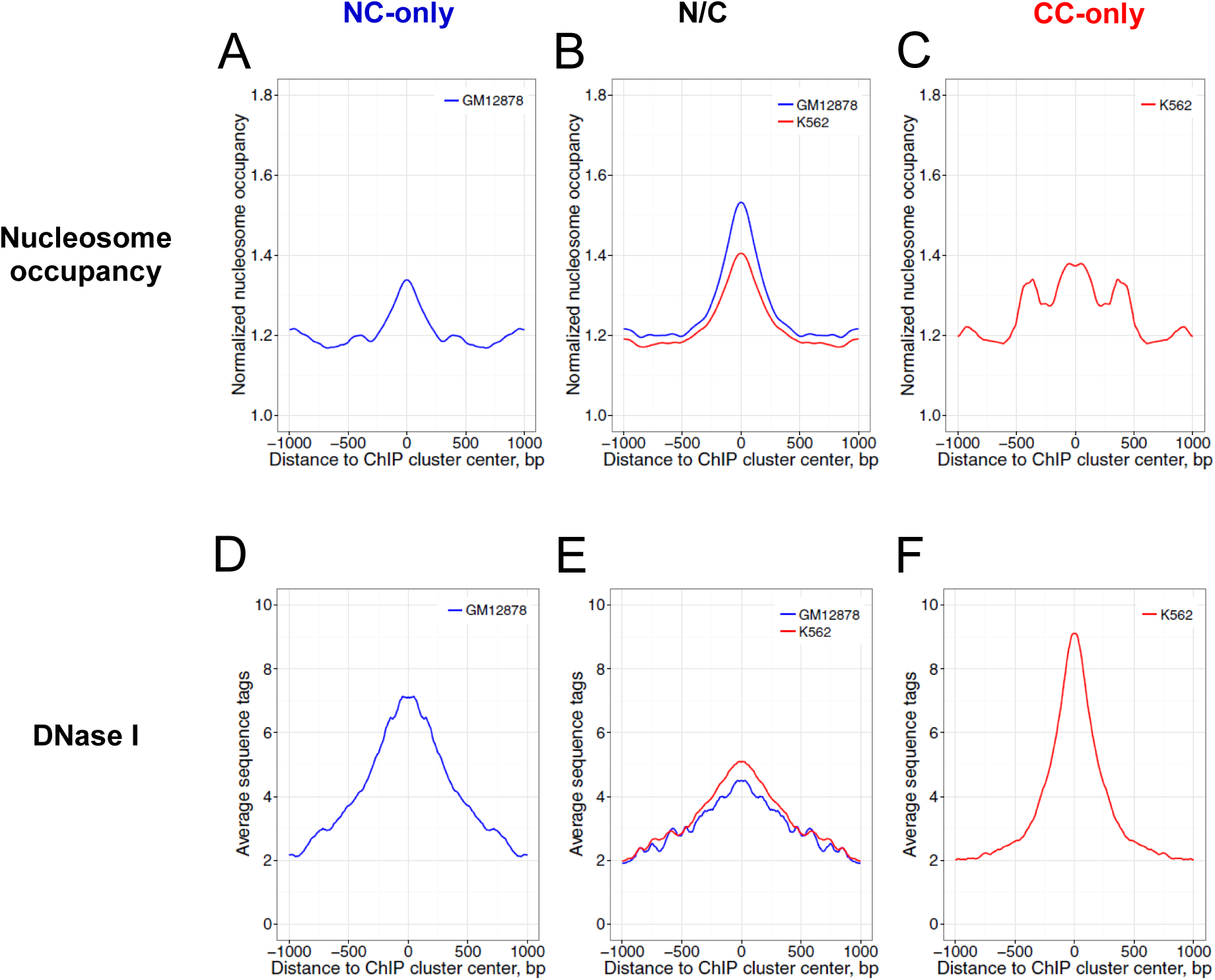
Profiles of nucleosome occupancy (A-C) and DNase I sensitivity (D-F) around p53 ChIP clusters of the NC group (A, D), the N/C group (B, E), and the CC group (C, F). All the ChIP clusters were aligned to their centers (position 0). Nucleosome mapping data and DNase-seq data for GM12878 cells (blue) and K562 cells (red) are shown. The nucleosome occupancy values were normalized with respect to the average value of the genome. The averaged nucleosome occupancy values and DNase-seq values were symmetrized with respect to the centers of the p53 ChIP clusters. The differences between position 0 and positions ±1000 in each group were evaluated statistically by Wilcoxon tests (Supplemental Table S6).

To examine the chromatin accessibility for NC and CC p53 ChIP clusters, we examined the profiles of DNase I sensitivity measured by DNase-seq in GM12878 and K562 chromatin (Fig. 4D-F). We observed that genomic regions near the p53 ChIP clusters exhibit increased DNase I sensitivity compared to flanking DNA (Wilcoxon test, *p* < 10^−15^), indicating that, in general, p53 ChIP clusters are located in accessible genomic regions.

In summary, we found that both NC and CC p53 ChIP clusters are characterized by two chromatin features. The first is high nucleosome occupancy, which is consistent with earlier reports on p53-induced DNA bending (Balagurumoorthy et al. 1995; Nagaich et al. 1997) and p53 interaction with its target sites exposed on the nucleosomal surface (Sahu et al. 2010; Cui et al. 2014). This nucleosome-binding ability distinguishes p53 from most TFs, which have functional DNA binding sites that are often found in nucleosome-depleted regions *in vivo* (Wang et al. 2012). The second feature is high DNase I sensitivity, which is shared by most TFs (Wang et al. 2012). Thus, unlike most TFs, p53 binding sites have a unique combination of characteristics, high nucleosome occupancy and high DNase I sensitivity.

### Epigenetic landscape around p53 ChIP clusters in normal and cancer cells

To further investigate the chromatin landscape around p53 ChIP clusters, we analyzed the histone modifications and DNA methylation data for GM12878 and K562 cells (Kundaje et al. 2012) which we downloaded from the ENCODE database. It is well established that lysine methylation of histone H3 and H4 is implicated in either transcriptional activation or repression, depending on the methylation sites. In particular, trimethylation of K4 and K36 on histone H3 (H3K4me3 and H3K36me3) are active histone marks. H3K4me3 is often found at gene promoters, whereas H3K36me3 is associated with transcribed regions in gene bodies (Greer and Shi 2012). By contrast, trimethylation of K27 (H3K27me3) is a repressive signal, primarily found at promoters in gene-rich regions.

We examined the epigenetic profiles of p53 ChIP clusters in the three groups and observed several interesting tendencies. First, NC p53 ChIP clusters have a higher level of transcriptionally active histone marks (H3K4me3 and H3K36me3) compared to their counterparts in cancer chromatin (Fig. 5 and Supplemental Fig. S11). In particular, the NC p53 ChIP clusters in GM12878 chromatin have a significantly higher level of H3K4me3 and H3K36 me3 than the CC ChIP clusters in K562 chromatin (compare Figs. 5A and 5C, and Figs. S11A and S11C). For the N/C group, the ChIP clusters in GM12878 chromatin also exhibit a higher level of active histone marks than in K562 chromatin (Fig. 5B and Supplemental Fig. S11B). Since H3K4me3 and H3K36me3 are often found in active genes (i.e., at promoter and gene body, respectively), our data strongly suggest that p53-DNA interactions in NC chromatin are associated with gene activation. In other words, p53-dependent transcriptional activation is more likely to occur in NC chromatin than in CC chromatin.

**Fig. 5.**
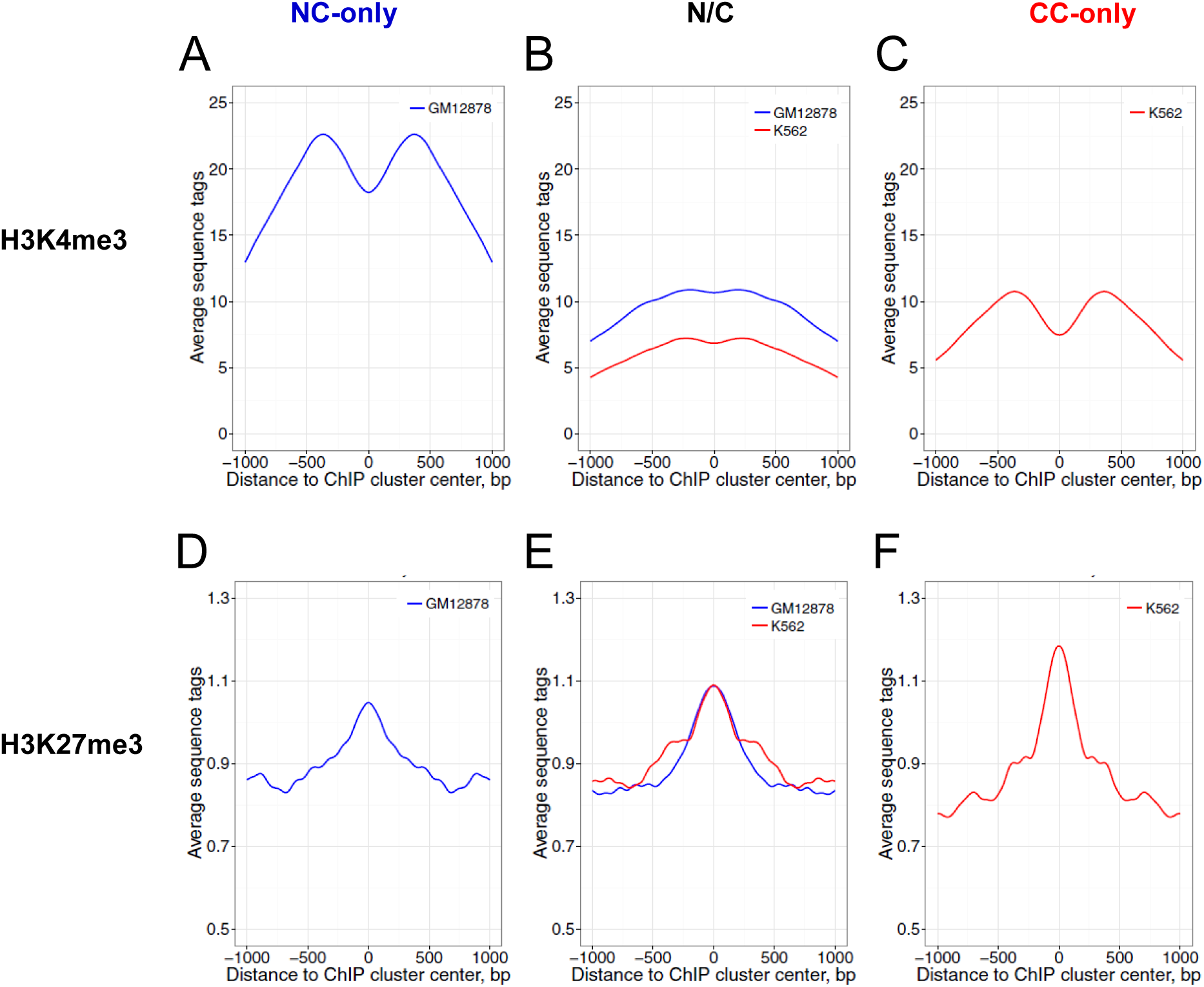
Profiles of the active histone mark H3K4me3 (A-C) and the repressive histone markH3K27me3 (D-F) around the p53 ChIP clusters in the NC-only group (A, D), the N/C group (B, E) and the CC-only group (C, F). The p53 ChIP clusters were aligned to their centers (position 0). The aggregate profiles of H3K4me3 and H3K27me3 data from GM12878 (blue) and K562 (red) are shown for the genomic region around the ChIP cluster centers. The averaged H3K4me3 and H3K27me3 values were symmetrized across the centers. The differences between various groups at position 0 were evaluated statistically by Wilcoxon tests (Supplemental Table S6).

Second, CC p53 ChIP clusters have a significantly higher level of repressive histone mark H3K27me3 than NC ChIP clusters (Figs. 5D-F). These data are consistent with the coupling of p53 binding in CC chromatin with transcriptional repression.

Third, the p53 ChIP clusters in the cancer-only group have a significantly lower level of DNA methylation than their counterparts in the NC-only group (Figs. 6A and 6C). Localization of the p53 CC-only sites in genomic regions with low DNA methylation may be related to chromatin decondensation due to global DNA hypomethylation in cancer. Unexpectedly, for the p53 ChIP clusters in the N/C group, DNA methylation in K562 chromatin is higher than that in GM12878 chromatin (Fig. 6B). Thus, in K562 cancer cell chromatin, the ChIP clusters in the N/C group exhibit a drastically different DNA methylation level from those in the CC-only group (Fig. 6B and 6C, Wilcoxon test, *p* = 8.0 × 10^−13^). These data suggest that the interplay between p53 binding and level of DNA methylation in CC chromatin is more complicated than that between p53 binding and histone H3 methylation.

**Fig. 6.**
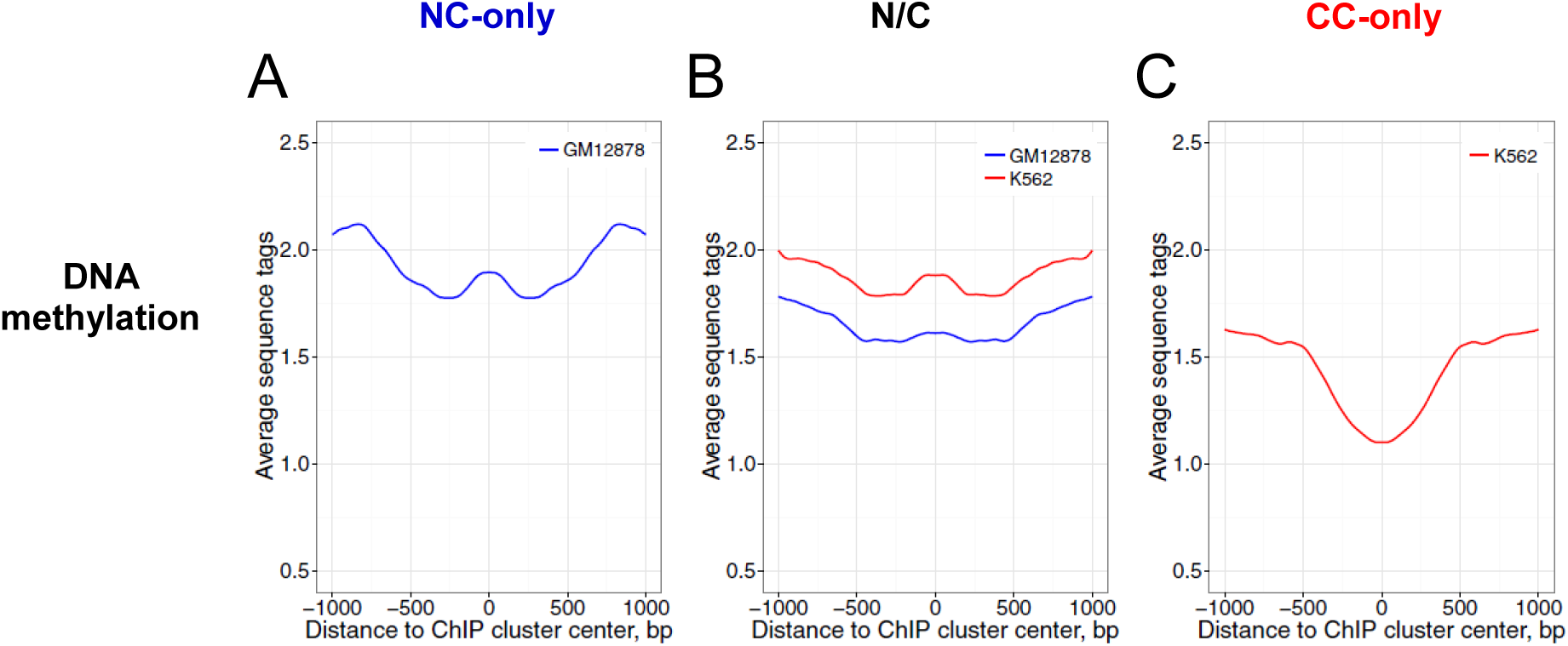
Profiles of DNA methylation around p53 ChIP cluster centers in the NC-only group (A), the N/C group (B) and the CC-only group (C). The MeDIP-seq data of the normal cell line GM12878 (blue) and the cancer cell line K562 (red) were used for this analysis. The MeDIP-seq data at genomic positions flanking the ChIP cluster centers were averaged and symmetrized across the centers (position 0). The differences between various groups at position 0 were evaluated statistically by Wilcoxon tests (Supplemental Table S6).

Note that our data differs from the findings of a previous study (Botcheva et al. 2011) in two aspects. First, Botcheva et al. reported that about 40% of p53 binding sites in NC chromatin (IMR90) are located <2 kb from TSSs, while only 15-20% of NC p53 sites were found in the same genomic regions in our study (Supplemental Fig. S7). Second, Botcheva et al. observed that p53 binding sites (in IMR90) are enriched in CpG islands and in hypomethylated DNA, perhaps due to their proximity to TSSs. Although it is highly probable that some NC ChIP clusters occur in hypomethylated DNA, our data indicate that overall, NC ChIP clusters tend to have a high level of DNA methylation (Fig. 6A). Note that the Botcheva data (for ~800 ChIP fragments) are included in our analysis and account for ~2% of the whole NC dataset (Supplemental Table S2). The observed differences are probably due to the methods used by Botcheva and colleagues to select the p53 binding sites from their ChIP-seq data.

## Discussion

### Over-representation of ERV1 family repeats in p53 high-confidence binding sites

As proposed in earlier studies (Britten and Davidson 1969), the transposable elements (TEs) are indispensable for evolution of the regulatory networks. Nowadays, this idea has been supported by the discovery of numerous repetitive elements exapted as cis-regulatory sequences (Bannert and Kurth 2004). It has been shown that species-specific TEs can greatly shape the transcriptional network of many TFs in humans and mice by contributing up to ~20% of functional binding sites (Bourque et al. 2008; Kunarso et al. 2010).

The tumor suppressor p53 is one of the TFs known to utilize TEs to expand its transcriptional networks. We and others have reported the existence of p53 binding sites in primate-specific interspersed repeats, including LTRs (Wang et al. 2007), SINEs such as Alu (Zemojtel et al. 2009; Cui et al. 2011) and LINEs (Harris et al. 2009). In particular, we have discovered that 15% of the functional p53 REs (24 out of 160) are located in human repeats, and more than half of these repeat-associated REs reside in Alu elements (Cui et al. 2011). Other studies (Wang et al. 2007; Zemojtel et al. 2009) focused on p53 ChIP fragments and found that 30-40% of ChIP fragments reside in repetitive regions. Note that only ~300 p53 ChIP fragments from a cancer cell line were analyzed in these studies, which only accounts for 0.25% of total fragments analyzed in this work (Supplemental Table S2). Our analysis showed that about 40% of p53 ChIP clusters occur in various classes of human repeats including SINE, LTR, LINE and simple repeats. We also found that p53 ChIP clusters are enriched in the ERV repeats, in agreement with the conclusion of previous studies (Wang et al. 2007). Although ERVs only comprise ~8% of the human genome (Lander et al. 2001), they account for 30-40% of p53 pile-12+ clusters (Fig. 3B-C). About 70-90% of the LTR-associated p53 motifs reside in ERV1 elements (Fig. 3E-F), highlighting an important role of the ERV1 repeat family in the p53 transcriptional program. Note that the ERV1 family is also over-represented in binding sites of pluripotency factors including OCT4 and NANOG in humans and mice (Kunarso et al. 2010). These findings suggest that ERVs act as important transcriptional regulatory elements that may be involved not only in developmental and tissue-specific gene regulation (Thompson et al. 2016), but also in tumor suppression.

It is generally believed that p53 negatively regulates TE transcription and inhibits retrotransposition to maintain genome stability. For example, it was observed that p53, together with DNA methylation, is involved in the epigenetic silencing of TEs (Leonova et al. 2012). In several other studies it was also found that p53 represses the transcription of HERV-1-LTRs (Chang et al. 2007) and Alu elements (Stein et al. 2002). However, in certain cellular contexts, p53 appears to increase the transcription of LINE-RNAs (Harris et al. 2009) and long intergenic non-coding RNAs (lincRNA) enriched in Alu repeats (Younger et al. 2015). These studies show that the functional significance of p53 occupancy in the repetitive regions is still far from being clear.

Finally, we found that p53 ChIP clusters with high PWM scores tend to overlap with human repeats (Fig. 2C and Fig. 3A-C). These data are consistent with previous findings (Su et al. 2015) that TE-associated p53 sites have a higher occupancy and correlate with stronger motif sequences, compared with non-TE-associated p53 binding sites. Su et al. described a strong correlation between occupancy and binding strength of p53 motifs (Su et al. 2015). In line with this view, our data suggest that the high occupancy of p53 in repetitive regions may be due to the existence of almost perfect p53 motifs in ERV LTRs (Wang et al. 2007) and Alu elements (Cui et al. 2011).

### High nucleosome occupancy and DNase I sensitivity: a characteristic chromatin feature of pioneer TFs including p53

The interaction between TFs and their cognate target sites is the central theme of gene regulation. In eukaryotes, DNA wraps around histone octamers to form nucleosomes (van Holde 1988), which often occlude TF binding. Hence, efficient binding of a given TF to its target site is greatly reduced in the context of chromatin (Owen-Hughes and Workman 1994). Therefore, a long-held notion is that TFs compete with nucleosomes to gain access to their binding sites (Workman and Kingston 1992). This notion was supported by the comprehensive analysis of ChIP fragments bound by 119 human TFs, which showed that on average the fragments are located in nucleosome-depleted, DNase I sensitive genomic regions (Wang et al. 2012)

However, recent studies have identified a group of TFs known as pioneer factors that can access their target sites in nucleosomal DNA (see review in Iwafuchi-Doi and Zaret 2014 in which 13 pioneer TFs are listed). Examples of pioneer TFs include FoxA family proteins (FoxA1, FoxA2 and FoxA3) and pluripotency factors (Oct4, Sox2 and Klf4). Earlier studies have suggested that p53-DNA binding also occurs in nucleosome-enriched regions. This idea first came from several observations that (i) p53 induces DNA bending when bound to its target (Balagurumoorthy et al. 1995), and (ii) the degree of p53-induced DNA bending correlates with binding affinity – the more pronounced the DNA bend, the higher the affinity of DNA interactions with p53 (Nagaich et al. 1997). Later, both experimental studies (Sahu et al. 2010; Laptenko et al. 2011) and computational modelling (Sahu et al. 2010) indicated that p53 can interact with its target sites on the surface of a nucleosome. These data are consistent with our observation that p53 binding sites are characterized with high nucleosome occupancy (Fig. 4). Therefore, p53 can be considered as one of the pioneer TFs (Iwafuchi-Doi and Zaret 2014; Sammons et al. 2015).

A previous study showed that the two well-known p53 REs in the *p21* gene reside in the DNase I-resistant domain (Braastad et al. 2003). By contrast, here we show that p53 ChIP clusters as a whole are in DNase I-sensitive regions (Fig. 4). Thus, p53 binding sites exhibit an unusual chromatin pattern: high nucleosome occupancy and high DNase I sensitivity, which distinguishes it from most DNA-binding TFs (Wang et al. 2012). Note that this unusual chromatin pattern is also observed for the binding sites of the pioneer factors PR (Ballare et al. 2013a; Ballare et al. 2013b) and FoxA2 (Iwafuchi-Doi et al. 2016). Therefore, we suggest that enhanced nucleosome occupancy and DNase I sensitivity may be a general feature of p53 and other pioneer TFs. This unique combination of chromatin features would allow access of pioneer TFs to their binding sites in chromatin prior to the access of other TFs.

### A new model for differential p53 binding to normal and cancer chromatin

We conducted a comprehensive analysis of p53 ChIP clusters from various NC and CC lines. Our work demonstrates that the p53 sites in NC and CC chromatin are characterized by distinct epigenetic features, highlighting the important role of the chromatin context in modulating p53 occupancy *in vivo* (Espinosa and Emerson, 2001; Shaked et al. 2008; Millau et al. 2010). In light of these findings, we propose a hypothetical model for distinctive p53 binding patterns in NCs and CCs (Fig. 7).

**Fig. 7.**
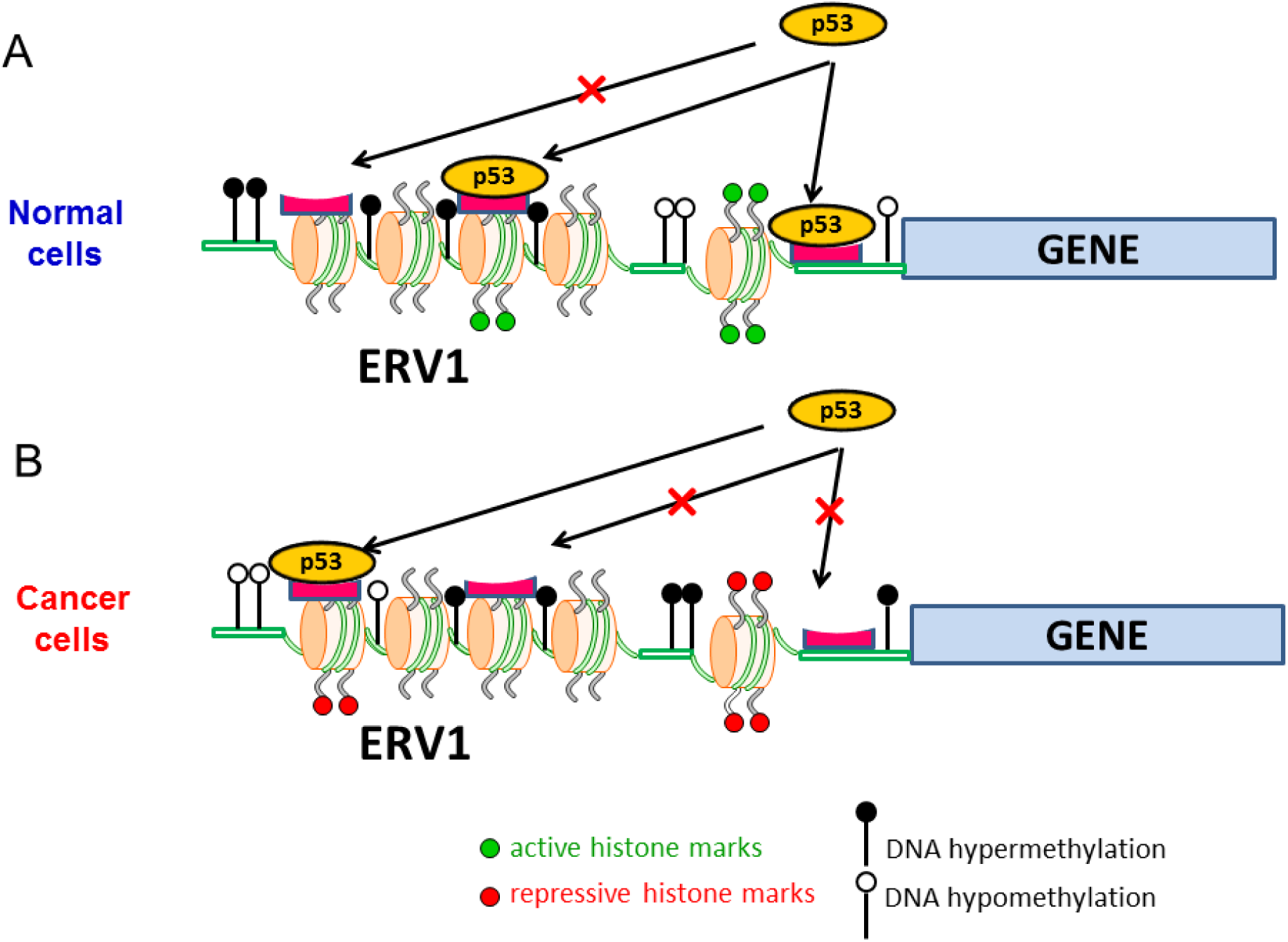
A model for p53 distinctive binding patterns in normal (A) and cancer (B) cells. In normal cells, p53 binding often occurs in genomic regions close to the gene TSS, characterized by high levels of DNA methylation (filled circles) and active histone marks such as H3K4me3 (green dots on histone tails). By contrast, the p53 binding sites in cancer cells are characterized by low levels of DNA methylation (empty circles) and high levels of repressive histone marks (red dots on histone tails). Reduction of DNA methylation levels leads to chromatin decondensation, which helps to expose p53 sites that are normally embedded in closed chromatin domains. Thus, the shift of the genomic binding patterns of p53 in cancer cells appears to reflect cancer-associated epigenetic dysregulation. Both normal and cancer p53 binding sites are enriched in ERV1 retroviral elements.

The NC p53 ChIP clusters are often found in genomic regions characterized by high levels of transcriptionally active histone marks such as H3K4me3 and H3K36me3 (Fig. 5). These data are consistent with earlier studies (Menendez et al. 2013) showing that both p53 binding and transactivation are associated with increased H3K4me3 levels. p53-dependent H3K4 methylation is mediated by histone-lysine N-methyltransferase SET1/MLL, which is enhanced by p53- and p300-dependent H3 acetylation (Tang et al. 2013). Note that H3K4me3 is often found in promoter regions, consistent with our observation that a substantial number (20-25%) of NC p53 binding sites are located <5 kb from a TSS (Supplemental Fig. S5).

By contrast, the CC p53 ChIP clusters are distinguished by a decreased level of active histone marks (H3K4me3 and H3K36me3) (Fig. 5) and an increased level of repressive histone marks (H3K27me3) (Supplemental Fig. S11). Moreover, the CC p53 sites are characterized by a lower level of DNA methylation, compared to their counterparts in normal cells (Fig. 6C), which may be due to genome-wide DNA hypomethylation in CC chromatin (Jones and Baylin 2007; Baylin and Jones 2011). We hypothesize that global DNA hypomethylation in cancer cells decondenses normally closed chromatin domains, which allows p53 to access new cognate sites in these domains. Taken together, these data suggest that many of these newly exposed p53 binding sites in CC chromatin fall within either genomic regions with repressive histone marks or “proto-enhancers” that are devoid of classic chromatin modifications (Sammons et al. 2015). This may explain why, on average, p53 ChIP clusters in CCs have low levels of active histone marks and high levels of repressive histone marks (Fig. 5). In either case, p53 can engage with its target sites in silent chromatin via pioneer factor activity and, in an appropriate cellular context, can recruit co-factors to activate the enhancers (Sammons et al. 2015). This pioneer activity requires the unique ability of p53 to interact with its target sites in a chromatin state that is non-permissive to most TFs.

Our study provides a simple clue to understanding the distinct p53 genomic binding patterns in normal and cancer cells. That is, p53 binding sites in NCs often occur in active chromatin characterized by H3K4me3 and H3K36me3 marks, which may be coupled with gene activation. By contrast, p53 binding sites in CCs are often found in repressive chromatin characterized by a high level of H3K27me3, which is likely to be associated with gene repression. This model could be tested by the analysis of gene expression data in normal and cancer cells. Understanding p53-dependent gene activation and repression in the context of NC and CC chromatin will shed new light on the complicated cellular mechanisms underlying p53 transcriptional regulation.

## Methods

### p53 response elements and ChIP fragments

The largest set of functional p53 REs so far has been collected (Riley et al. 2008), and comprises 156 binding sites. Note, however, that a p53 site from human hepatitis B virus (HBV) and the overlapping sites from the *CDKN1A/p21* gene were included in this data. We removed the HBV site and one of the *p21* sites. The final list contains 154 REs. Based on the sequences of the p53 REs, we identified their genomic locations in the hg18 human reference genome (Supplemental Table S1).

A total of 25 datasets of p53 ChIP fragments were collected (Supplemental Table S2), including 9 from NC lines and 16 from CC lines. The locations of the fragments were downloaded from the papers and, if needed, ‘converted’ to corresponding locations in the human genome assembly hg18 using the LiftOver utility at the University of California Santa Cruz (UCSC) Genome Browser web server.

Note that only genomic binding data sets mapped with endogenous wild-type p53 were selected in this study. Those data sets with p53 mutants (Do et al. 2012; Schlereth et al. 2013; Wang et al. 2014), family members (Yang et al. 2006; Yang et al. 2010) or p53 in other mammals (Li et al. 2012) were not used.

### Non-ENCODE nucleosome data sets

Nucleosomes were reconstituted with human granulocyte DNA and recombinantly derived histone octamers to produce a genome-wide *in vitro* nucleosome map (Valouev et al. 2011). The raw sequence reads from these nucleosomes were downloaded from the NCBI Gene Expression Omnibus (GSE25133). The reads were mapped to human genome assembly hg18 using the Bowtie aligner with the default settings. Only the reads that were uniquely mapped to the genome were used and these reads were extended to 147 bp in the 5’ to 3’ direction. Normalization of nucleosome occupancy across the genome was performed as described before (Cui et al. 2012). Briefly, for each nucleotide position in the genome, the total number of nucleosomal DNA sequences covering this position was divided by the average number of nucleosomal sequences per base pair across the genome. The resulting value was assigned to this position as the normalized nucleosome occupancy value.

### ENCODE MNase-seq, ChIP-seq, MeDIP-seq, and DNase-seq data sets

The MNase-seq reads for in vivo nucleosomes in both GM12878 and K562 cell lines were downloaded from the UCSC Genome Browser-FTP server (ftp://hgdownload.cse.ucsc.edu/goldenPath/hg19/encodeDCC/wgEncodeSydhNsome/). The BAM files in the human genome assembly hg19 were downloaded. The BAM files were then converted to SAM files using SAMtools (Li et al. 2009). The reads were then extended to 147 bp in the 5’ to 3’ direction, and the normalized nucleosome occupancy at each position was calculated, as described before (Cui et al. 2012). The normalized values at each position were smoothed with a 60-bp window.

The ChIP-seq data for histone marks H3K4me3, H3K27me3 and H3K36me3 from both GM12878 and K562 cell lines were downloaded from the UCSC Genome Browser web server (http://genome.ucsc.edu/cgi-bin/hgFileUi?db=hg19&g=wgEncodeUwHistone). The BAM files in human genome assembly hg19 were downloaded. The BAM files were then converted to SAM files using SAMtools. The start position of each read was shifted by 73 bp in the 5’ to 3’ direction. The values at each position were smoothed twice with a 60-bp window.

DNA methylation MeDIP-seq data and DNase-seq data from both GM12878 and K562 cell lines were downloaded from NCBI GEO datasets (GSM1368906, GSM1368907, GSM816655 and GSM816665). The BigWig files in human genome assembly hg19 were downloaded. The values at each position were smoothed with a 60-bp window.

Since the p53 ChIP clusters are in human genome assembly hg18, the UCSC LiftOver utility was used along with the hg18-hg19 chain file to convert the sites to their corresponding positions in human genome assembly hg19 to make the profiles.

### Monte Carlo simulation

A Monte Carlo simulation was performed to assess the background level of overlapping ChIP fragments obtained from published studies (Supplemental Table S2) using BEDTools (Quinlan and Hall 2010). In the simulation, 42,020 genomic DNA segments (1290 bp on average) and 77,796 segments (663 bp on average) were randomly selected from human genome assembly hg18, and the number of fragments overlapped with others was determined. This process was repeated 100 times to compute the percentage of randomly selected DNA fragments that overlapped. The results are shown in Supplemental Table S3. For NC ChIP fragments, we estimated that about 11% of pile-2 clusters, 0.5% of pile-3 clusters and < 0.001% of pile-4 clusters can be formed by random sampling. Similar results were obtained for CC ChIP fragments. This suggests that about 88% of the pile-2 and over 99.5% of pile-3 or higher (denoted as pile-3+) clusters likely represent real ChIP enrichment events in normal or cancer cells. Because the pile-2 clusters contain a substantial number of false positive clusters (~12%), we selected pile-3+ clusters for further analysis.

### Repeat analysis

Human repetitive region positions in human genome assembly hg18 were downloaded from the UCSC Genome Browser. The repeat elements were identified using RepeatMasker (v3.2.7) and Repbase Update (9.11). The major types of repeat elements were selected for analysis, including SINE (Alu, MIR), LINE (CR1, L1, L2 and RTE), LTR (ERV1, ERVK, ERVL and Gypsy), Simple Repeat ((TG)_n_, (TCG)_n_, (CACTC)_n_, (GAGTG)_n_, and (TATATG)_n_), Low Complexity (C-rich, GC-rich, GA-rich, CT-rich), DNA (MuDR, PiggyBac, TcMar-Mariner, hAT-Charlie). The remaining repeat types were included into the “Other” category.

A p53 motif was classified to be in a repeat (sub)family if it overlaps with any repeat element in that (sub)family. The p53 motifs that are not covered by any repeat elements are classified as a “Non_repeat” group.

### Functional Annotation

The enriched biological pathways were identified by DAVID 6.7 (Huang et al. 2009a; Huang et al. 2009b) using the genes associated with p53 ChIP clusters. If the TSS of a gene was located within 5 kb from the center of a ChIP cluster, this gene was selected for the analysis. Most enriched pathways were determined using DAVID Annotation Chart Analysis and the Kyoto Encyclopedia of Genes and Genomes (KEGG) database. All genes associated with the ChIP clusters in the NC-only, N/C and CC-only groups were analyzed by DAVID as a group.

## Acknowledgements

The authors are grateful to G. Leiman for editing the text. The research was supported by a NIH grant R15GM116102 (to F. C.) and The Intramural Research Program of the National Institutes of Health (Center for Cancer Research, National Cancer Institute) (to V. B. Z.). Funding for open access charge: NIH grant R15GM116102.

## References

Akdemir KC, Jain AK, Alton K, Aronow B, Xu X, Cooney AJ, Li W, Barton MC. 2014. Genome-wide profiling reveals stimulus-specific functions of p53 during differentiation and DNA damage of human embryonic stem cells. Nucleic Acids Res. 42: 205–223.

Balagurumoorthy P, Sakamoto H, Lewis MS, Zambrano N, Clore GM, Gronenborn AM, Appella E, Harrington RE. 1995. Four p53 DNA-binding domain peptides bind natural p53 response elements and bend the DNA. Proc. Natl. Acad. Sci. USA 92: 8591–8595.

Ballare C, Castellano G, Gaveglia L, Althammer S, Gonzalez-Vallinas J, Eyras E, Le Dily F, Zaurin R, Soronellas D, Vicent GP, Beato M. 2013a. Nucleosome-driven transcription factor binding and gene regulation. Mol. Cell 49: 67–79.

Ballare C, Zaurin R, Vincent GP, Beato M. 2013b. More help than hindrance. Nucleus 4: 189–194.

Bannert N, Kurth R. 2004. Retroelements and the human genome: new perspectives on an old relation. Proc. Natl. Acad, Sci. USA 101: 14572–14579.

Barlev NA, Liu L, Chehab NH, Mansfield K, Harris KG, Halazonetis TD, Berger SL. 2001. Acetylation of p53 activates transcription through recruitment of coactivators/histone acetyltransferases. Mol. Cell 8: 1243–1254.

Baylin SB, Jones PA. 2011. A decade of exploring the cancer epigenome – biological and translational implications. Nat. Rev. Cancer 11: 726–734.

Beckeman R, Prives C. 2010. Transcriptional regulation by p53. Cold Spring Harb. Perspect. Biol. 2: a000935.

Botcheva K, McCorkle SR, McCombie WR, Dunn JJ, Anderson CW. 2011. Distinct p53 genomic binding patterns in normal and cancer-derived human cells. Cell Cycle 10: 4237–4249.

Botcheva K, McCorkle SR. 2014. Cell context dependent p53 genome-wide binding patterns and enrichment at repeats. PLoS One 9: e113492.

Bourque G, Leong B, Vega VB, Chen X, Lee YL, Srinivasan KG, Chew JL, Ruan Y, Wei CL, Ng HH, Liu ET. 2008. Evolution of the mammalian transcription factor binding repertoire via transposable elements. Genome Res. 18: 1752–1762.

Boyle AP, Davis S, Shulha HP, Meltzer P, Margulies EH, Weng Z, Furey TS, Crawford GE. 2008. High-resolution mapping and characterization of open chromatin across the genome. Cell 132: 311–322.

Braastad CD, Han Z, Hendrickson EA. 2003. Constitutive DNase I hypersensitivity of p53-regulated promoters. J. Biol. Chem. 278: 8261–8268.

Britten RJ, Davidson EH. 1969. Gene regulation for higher cells: a theory. Science 165: 349–357.

Candau R, Scolnick DM, Darpino P, Ying CY, Halazonetis TD, Berger SL. 1997. Two tandem and independent subactivation domains in the amino terminus of p53 require the adaptor complex for activity. Oncogene 15: 807–816.

Chang NT, Yang WK, Huang HC, Yeh KW, Wu CW. 2007. The transcriptional activity of HERV-I LTR is negatively regulated by its cis-elements and wild type p53 tumor suppressor protein. J. Biomed. Sci. 14: 211–222.

Contente A, Dittmer A, Koch MC, Roth J, Dobbelstein M. 2002. A polymorphic microsatellite that mediates induction of PIG3 by p53. Nat. Genet. 30: 315–320.

Cui F, Sirotin MV, Zhurkin VB. 2011. Impact of Alu repeats on the evolution of human p53 binding sites. Biol. Direct 6: 2.

Cui F, Cole HA, Clark DJ, Zhurkin VB. 2012. Transcriptional activation of yeast genes disrupts intragenic nucleosome phasing. Nucleic Acids Res. 40: 10753–10764.

Cui F, Zhurkin VB. 2014. Rotational positioning of nucleosomes facilitates selective binding of p53 to response elements associated with cell cycle arrest. Nucleic Acids Res. 42: 836–847.

Do PH, Varanasi L, Fan S, Li C, Kubucka I, Newman V, Chauhan K, Daniels SR, Boccetta M, Garrett MR, Li R, Martinez LA. 2012. Mutant p53 cooperates with ETS2 to promote etoposide resistance. Genes Dev. 26: 830–845.

el-Deiry WS, Kern SE, Pietenpol JA, Kinzler KW, Vogelstein B. 1992. Definition of a consensus binding site for p53. Nat. Genet. 1: 45–49.

el-Deiry WS, Tokino T, Velculescu VE, Levy DB, Parsons R, Trent JM, Lin D, Mercer WE, Kinzler KW, Vogelstein B. 1993. WAF1, a potential mediator of p53 tumor suppression. Cell 75: 817–825.

Espinosa JM, Emerson BM. 2001. Transcriptional regulation by p53 through intrinsic DNA/chromatin binding and site-directed cofactor recruitment. Mol. Cell 8: 57–69.

Fischer M, Quaas M, Nickel A, Engeland K. 2015. Indirect p53-dependent transcriptional repression of Survivin, CDC25C and PLK1 genes requires the cyclin-dependent kinase inhibitor p21/CDKN1A and CDE/CHR promoter sites binding the DREAM complex. Oncotarget 6: 41402–41417.

Gohler T, Reimann M, Cherny D, Walter K, Warnecke G, Kim E, Deppert W. 2002. Specific interaction of p53 with target binding sites is determined by DNA conformation and is regulated by the C-terminal domain. J. Biol. Chem. 277: 41192–41203.

Greer EL, Shi Y. 2012. Histone methylation: a dynamic mark in health, disease and inheritance. Nat. Rev. Genet. 13: 343–357.

Harris CR, Dewan A, Zupnick A, Normart R, Gabriel A, Prives C, Levine AJ, Hoh J. 2009. p53 response elements in human retrotransposons. Oncogene 28: 3857–3865.

Huang DW, Sherman BT, Lempicki RA. 2009a. Systematic and integrative analysis of large gene lists using DAVID Bioinformatics Resources. Nature Protoc. 4: 44–57.

Huang DW, Sherman BT, Lempicki RA. 2009b. Bioinformatics enrichment tools: paths toward the comprehensive functional analysis of large gene lists. Nucleic Acids Res. 37: 1–13.

Iwafuchi-Doi M, Zaret KS. 2014. Pioneer transcription factors in cell reprogramming. Genes Dev. 28: 2679–2692.

Iwafuchi-Doi M, Donahue G, Kakumanu A, Watts JA, Mahony S, Pugh BF, Lee D, Kaestner KH, Zaret KS. 2016. The pioneer transcription factor FoxA maintains an accessible nucleosome configuration at enhancers for tissue-specific gene activation. Mol. Cell 62: 79–91.

Johnson DS, Mortazavi A, Myers RM, Wold B. 2007. Genome-wide mapping of in vivo protein-DNA interactions. Science 316: 1497–1502.

Jones PA, Baylin SB. 2007. The epigenomics of cancer. Cell 128: 683–692.

Kaneshiro K, Tsutsumi S, Tsuji S, Shirahige K, Aburatani H. 2007. An integrated map of p53-binding sites and histone modification in the human ENCODE regions. Genomics 89: 178–188.

Kunarso G, Chia NY, Jeyakani J, Hwang C, Liu X, Chan YS, Ng HH, Bourque G. 2010. Transposable elements have rewired the core regulatory network of human embryonic stem cells. Nat. Genet. 42: 631–634.

Kundaje A, Kyriazopoulou-Panagiotopoulou S, Libbrecht M, Smith CL, Raha D, Winters EE, Johnson SM, Snyder M, Batzoglou S, Sidow A. 2012. Ubiquitous heterogeneity and asymmetry of the chromatin environment at regulatory elements. Genome Res. 22: 1735–1747.

Lander ES, Linton LM, Birren B, Nusbaum C, Zody MC, Baldwin J, Devon K, Dewar K, Doyle M, FitzHugh Wet al.2001. Initial sequencing and analysis of the human genome. Nature 409: 860–921.

Laptenko O, Beckerman R, Freulich E, Prives C. 2011. p53 binding to nucleosomes within the p21 promoter in vivo leads to nucleosome loss and transcriptional activation. Proc. Natl. Acad. Sci. USA 108: 10385–10390.

Leonova KL, Brodsky L, Lipchick B, Pal M, Novototskaya L, Chenchik AA, Sen GC, Komarova EA, Gudkov AV. 2012. p53 cooperates with DNA methylation and a suicidal interferon response to maintain epigenetic silencing of repeats and noncoding RNAs. Proc. Natl. Acad. Sci. USA 110: E89–E98.

Levine AJ. 1997. p53, the cellular gatekeeper for growth and division. Cell 88: 323–331.

Li H, Handsaker B, Wysoker A, Fennell T, Ruan J, Homer N, Marth G, Abecasis G, Durbin R, and 1000 Genome Project Data Processing Subgroup. 2009. The sequence alignment/map (SAM) format and SAMtools. Bioinformatics 25: 2078–2079.

Li M, He Y, Dubois W, Wu X, Shi J, Huang J. 2012. Distinct regulatory mechanisms and functions for p53-activated and p53-repressed DNA damage response genes in embryonic stem cells. Mol. Cell 46: 30–42.

Lidor NE, Field Y, Lubling Y, Widom J, Oren M, Segal E. 2010. p53 binds preferentially to genomic regions with high DNA-encoded nucleosome occupancy. Genome Res. 20: 1361–1368.

McDade SS, Patel D, Moran M, Campbell J, Fenwick K, Kozarewa I, Orr NJ, Lord CJ, Ashworth AA, McCance DJ. 2014. Genome-wide characterization reveals complex interplay between TP53 and TP63 in response to genotoxic stress. Nucleic Acids Res. 42: 6270–6285.

Menendez D, Nguyen T, Freudenberg JM, Mathew VJ, Anderson CW, Jothi R, Resnick M. 2013. Diverse stresses dramatically alter genome-wide p53 binding and transactivation landscape in human cancer cells. Nucleic Acids Res. 41: 7286–7301.

Millau JF, Bandele OJ, Perron J, Bastien N, Bouchard EF, Gaudreau L, Bell DA, Drouin R. 2010. Formation of stress-specific p53 binding patterns is influenced by chromatin but not by modulation of p53 binding affinity to response elements. Nucleic Acids Res. 39: 3053–3063.

Nagaich CK, Appella E, Harrington RE. 1997. DNA bending is essential for the site-specific recognition of DNA response elements by the DNA binding domain of the tumor suppressor protein p53. J. Biol. Chem. 272: 14842–14849.

Nozell S, Chen X. 2002. p21B, a variant of p21(Waf1/Cip1), is induced by the p53 family. Oncogene 21: 1285–1294.

Owen-Hughes T, Workman JL. 1994. Experimental analysis of chromatin function in transcription control. Crit. Rev. Eukaryot. Gene Exp. 4: 403–441.

Quinlan AR, Hall IM. 2010. BEDTools: a flexible suite of utilities for comparing genomic features. Bioinformatics 26: 841–842.

Riley T, Sontag E, Chen P, Levine A. 2008. Transcriptional control of human p53-regulated genes. Nat. Rev. Mol. Cell Biol. 9: 402–412.

Sahu G, Wang D, Chen CB, Zhurkin VB, Harrington RE, Appella E, Hager GL, Nagaich AK. 2010. p53 binding to nucleosomal DNA depends on the rotational positioning of DNA response element. J. Biol. Chem. 285: 1321–1332.

Sammons MA, Zhu J, Drake AM, Berger SL. 2015. TP53 engagement with the genome occurs in distinct local chromatin environments via pioneer factor activity. Genome Res. 25: 179–188.

Schlereth K, Heyl C, Krampitz AM, Mernberger M, finkernagel F, Scharfe M, Jarek M, Leich E, Rosenwald A, Stiewe T. 2013. Characterization of the p53 cistrome – DNA binding cooperativity dissects p53’s tumor suppressor functions. PLoS Genet. 9: e1003726.

Shaked H, Shiff I, Kott-Gutkowski M, Siegfried Z, Haupt Y, Simon I. 2008. Chromatin immunoprecipitation-on-chip reveals stress-dependent p53 occupancy in primary normal cells but not in established cell lines. Cancer Res. 68: 9671–9677.

Smeenk L, van Heeringen SJ, Koeppel M, van Driel MA, Bartels SJ, Akkers RC, Denissov S, Stunnenberg HG, Lohrum M. 2008. Characterization of genome-wide p53-binding sites upon stress response. Nucleic Acids Res. 36: 3639–3654.

Stein T, Crighton D, Warnock LJ, Milner J, White RJ. 2002. Several regions of p53 are involved in repression of RNA polymerase III transcription. Oncogene 21: 5540–5547.

Su D, Wang X, Campbell MR, Song L, Safi A, Crawford GE, Bell DA. 2015. Interactions of chromatin context, binding site sequence content, and sequence evolution in stress-induced p53 occupancy. PLoS Genet. 11: 1004885.

Thompson PJ, Macfaclan TS, Lorincz MC. 2016. Long terminal repeats: from parasitic elements to building blocks of the transcriptional regulatory repertoire. Mol. Cell 62: 766–776.

Tang Z, Chen WY, Shimada M, Nguyen UT, Kim J, Sun XJ, Sengoku T, McGinty RK, Fernandez JP, Muir TW, Roeder RG. 2013. SET1 and p300 act synergistically, through coupled histone modifications, in transcriptional activation by p53. Cell 154: 297–310.

Valouev A, Johnson SM, Boyd SD, Smith CL, Fire AZ, Sidow A. 2011. Determinants of nucleosome organization in primary human cells. Nature 474: 516-520. van Holde KE. 1988. Chromatin. Springer-Verlag, New York.

Vrba L, Junk DJ, Novak P, Futscher BW. 2008. p53 induces distinct epigenetic states at its direct target promoters. BMC Genomics 9: 486.

Wang T, Zeng J, Lowe CB, Sellers RG, Salama SR, Yang M, Burgess SM, Brachmann RK, Haussler D. 2007. Species-specific endogenous retroviruses shape the transcriptional network of the human tumor suppressor protein p53. Proc. Natl. Acad. Sci. USA 104: 18613–18618.

Wang J, Zhuang J, Iyer S, Lin X, Whitfield TW, Greven MC, Pierce BG, Dong X, Kundaje A, Cheng Y. et al. 2012. Sequence features and chromatin structure around the genomic regions bound by 119 human transcription factors. Genome Res. 22: 1798–1812.

Wang B, Niu D, Lam TH, Xiao Z, Ren EC. 2014. Mapping the p53 transcriptome universe using p53 natural polymorphs. Cell Death Diff. 21: 521–532.

Wei CL, Wu Q, Vega VB, Chiu KP, Ng P, Zhang T, Shahab A, Yong HC, Fu Y, Weng Z. et al. 2006. A global map of p53 transcription-factor binding sites in the human genome. Cell 124: 207–219.

Workman JL, Kingston RE. 1992. Nucleosome core displacement in vitro via a metastable transcription factor-nucleosome complex. Science 258: 1780–1784.

Yang A, Zhu Z, Kapranov P, McKeon F, Church GM, Gingeras TR, Struhl K. 2006. Relationships between p63 binding, DNA sequence, transcription activity, and biological function in human cells. Mol. Cell 24: 593–602.

Yang A, Zhu Z, Kettenbach A, Kapranov P, McKeon F, Gingeras TR, Struhl K. 2010. Genome-wide mapping indicates that p73 and p63 co-occupy target sites and have similar DNA-binding profiles in vivo. PLoS One 5: e11572.

Younger ST, Kenzelmann-Broz D, Jung H, Attardi LD, Rinn JL. 2015. Integrative genomic analysis reveals widespread enhancer regulation by p53 in response to DNA damage. Nuclear Acids Res. 43: 4447–4462.

Yu J, Zhang L, Hwang PM, Kinzler KW, Vogelstein B. 2001. PUMA induces the rapid apoptosis of colorectal cancer cells. Mol. Cell 7: 673–682.

Zemojtel T, Kielbasa SM, Arndt PF, Chung H-R, Vingron M. 2009. Methylation and deamination of CpGs generate p53-binding sites on a genomic scale. Trends Genet. 25: 63–66.

